# Enhanced effective codon numbers to understand codon usage bias

**DOI:** 10.1101/644609

**Authors:** Reginald Smith

## Abstract

Codon usage bias is a well recognized phenomenon but the relative influence of its major causes: G+C content, mutational biases, and selection, are often difficult to disentangle. This paper presents methods to calculate modified effective codon numbers that allow the investigation of the sources of codon bias and how genes or organisms have their codon biases shaped. In particular, it demonstrates that variation in codon usage bias across organisms is likely driven more by likely mutational forces while the variation in codon usage bias within genomes is likely driven by codon selectional forces.

**Author summary:** A new method of disaggregating codon bias influences is described where I show how that different values of the effective codon number, following Wright’s *N_c_*, can be used as ratios to demonstrate the similar or different causes of codon biases across genes or organisms. By calculating ratios of the different types of effective codon numbers, one can easily compare organisms or different genes while controlling for gene G+C content or codon nucleotide G+C content. The driving forces determining the variations in codon usage bias across or within organisms thus become much clearer.

## Introduction

From the decipherment of the genetic code [1] to early predictions of selection against supposedly neutral synonymous codons [2], the phenomenon of codon usage bias, the uneven usage of synonymous codons for amino acids [3,4], has been found to be ubiquitous not only across different organisms but even across different genes within a genome with those more highly expressed genes most likely to have codon biases [5–9]. Changes are driven by a variety of processes but they fall under two broad mechanisms: mutational biases which alter the codons, particularly via the nucleotide in the third position of a codon, in a manner that biases the codon frequency and the forces of selection which favor certain synonymous codons, often due to advantages such as translation efficiency [9–12]. Codon usage bias can also be driven by genome or gene G+C content where codons with higher G+C content are more prevalent under the influence of certain processes, for example the GC-biased gene conversion (gBGC) process in meiotic repair, that prefer G and C bases. Isochores within the genome can have varying G+C content which is relatively homogenous within the isochore but differs between isochores. Codon bias can differ between these reasons solely due to G+C content as well.

In practice, however, differentiating the effects of mutation or GC biased processes and selection on codon bias can be difficult. Different genes and even different locations on genes can have varying G+C content and different impacts from mutation processes. This can make G+C content and expected codon bias differ, even where mutation caused bias and selection are not prominent. To somewhat clarify these questions, we will take an approach that focuses on how different types of processes are relatively selectively neutral for certain groups of synonymous codons. Therefore, rather than try to exactly narrow effects to “mutation” or “selection”, we can describe how different aspects of codon bias are implicitly invariant amongst certain groups of synonymous codons.

Critical to the understanding of the underlying causes of codon usage bias has been the metrics used to define and measure it. This paper will supplement the most commonly used metric, Wright’s *N_c_* [13], hereafter designated as *N_c_*. First, we will briefly review the most common codon usage bias metrics and their particular advantages. Second, we will explain combinatorics using information theory and show how this can re-derive several *N_c_* like quantities that represent the different effects of genome or gene G+C content, codon nucleotide G+C content, and selection on specific codon usage. Finally, we will demonstrate the metrics’ utilization both across a wide group of organisms and the genes of several organisms to demonstrate how to measure the relative effects of biased mutation and selection in shaping codon usage bias.

### 0.1 Measurements of codon usage bias

From the beginning, various numerical metrics have been proposed in order to understand codon usage bias. Early measures used the relative frequency of synonymous codons against a maximum frequency within the same group to calculate codon usage bias. Metrics such as the relative synonymous codon usage index (RSCU) [9] and the codon adaptation index (CAI) [14] measured the usage of synonymous codons against random or maximum frequency focusing on measuring the relative disparity within the code of each amino acid. Later, and probably most prominent, was the work of Wright [13] whose effective number of codons, *N_c_*, used concepts of minimum homozygosity and the effective population size (considering each synonymous codon as an ‘allele’) to estimate codon usage bias. *N_c_* is one of the most widely used metrics and most useful for shorter genes though its value can exceed the actual numbers of codons in use. It has a maximum of 61 (64 total codons minus 3 stop codons in the standard code) and a minimum of 20 (one codon per amino acid). There have been several adaptations and commentaries on *N_c_* when amino acids are missing or exist only at low frequencies [15–18]. Similar to this paper, many codon usage measurements have also implemented information theoretic methods such as entropy in order to analyze codon usage bias. Amongst the first was Tavare and Song [19]. Zeeberg [20] calculated the information entropy in bits across synonymous codons and compared it to the G+C content in different codon positions across genes for the newly sequenced human and mouse genomes. Later, a new metric, synonymous codon usage order (SCUO) [21] also used information entropy but used the proportion of theoretical maximum entropy to create a metric demonstrating the relative diversity of codon usage from a value of 0 representing maximum diversity (random usage) up to 1 for extremely skewed codon usage. A measure of relative entropy [22] also was developed.

While many of these codon usage metrics have their own particular advantages such as easily interpretable values, dealing with extreme bias cases, etc. most works still demonstrate that the traditional *N_c_* and its variants perform reasonably well in comparison [23,24]. Therefore, a technique that can use the power of information theory as well as the general utility of *N_c_* can provide insight while combining the strengths of both.

## Materials and methods

### Combinatorics of codon bias

It is well known that for a nucleotide sequence of length *L*, there are at most, 4*^L^* possible different sequences using each nucleotide under the assumption they occur with equal frequency individually and relative to other nucleotides. Even for short sequences, the number of combinations soon becomes astronomical. However, such a sequence structure is essentially random while the sequences of living organisms are not [25].

A more constrained measure of the number of possible sequences that takes into account differing frequencies of occurrence uses the entropy function. Shannon and Weaver [26] showed that given a sequence consisting of *M* distinct symbols, where the frequency of the kth symbol is given by *p_k_*, the entropy *H* (in bits) is measured by

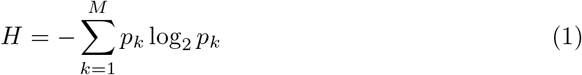

The expected number of sequences with symbol entropy *H* and of symbol length *L*, designated *N*, is given by

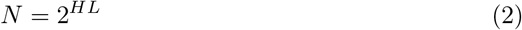

The entropy function represents not the information contained in any given sequence per se but how much a reduction in uncertainty (information) the sequence conveys given the frequency of occurrence in its symbols. This is easily applied to the nucleotide sequence case. For the four nucleotides, the Eq. 1 is accurate given the entropy calculated by the frequency of each base. When all occur equally, *H* = 2 and we get the original result 4*^L^*. A brief example of this technique is illustrative.

Assume that the base pairs G/C or A/T occur in equal combinations within a sequence and the G+C content proportion is given by *p_GC_*. The sequence entropy is then

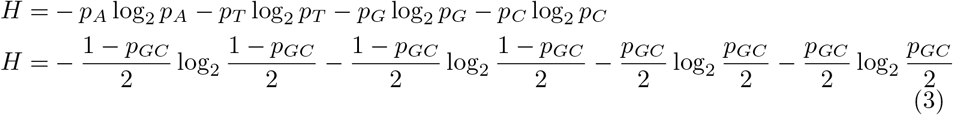

Therefore, the G+C content of the sequence can allow us to determine its entropy where

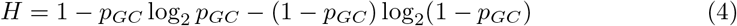

Based on this we can see how the change in G+C content alone can drastically reduce the number of possible sequences. A sequence of 100 bp with a G+C content of 50% will have *H* = 2 and 2^100×2^ expected combinations. For a G+C content of 60%, *H* = 1.97 and the sequence will have 2^100×1.97^ combinations. The ratio of this to the 50% case shows there are only 13% (2^100×.03^) as many expected combinations as in the case where G+C is 50%. The difference becomes more stark the longer the sequence under investigation becomes. At 1 kbp, a sequence of G+C at 60% will only have about 10^-9^ times as many combination as in the G+C = 50% case. While these are huge reductions, they still leave a large number of sequences possible.

One of the key questions in codon usage bias is the relative importance of factors such as G+C content, biased mutation rates, and selection in determining the usage pattern of synonymous codons. All factors play a part though it is well known that codon usage bias tends to correlate with levels of gene expression given different synonymous codons confer efficiency to the protein translation process. While it can be difficult to analyze each of these factors separately, one approach is to compare measures of codon bias in actual data with their expected values if only one or two of these factors alone skewed synonymous codon usage.

Using *N_c_* as our measurement of codon bias, we can derive alternate versions of *N_c_* which are due to primarily genome or gene wide G+C content, codon nucleotide G+C preference, which is often interpreted as a possible sign of mutation-caused bias, and codon selection processes. By comparing these to each other as well as the traditional definition of *N_c_* we can illuminate for individual organisms or even large groups of related organisms, how various processes shape codon usage bias.

## 1 Alternative measures of *N_c_*

We will define four types of additional *N_c_* as detailed in Table 1. All definitions will include only sense codons and exclude the three stop codons in the standard code. The first, *N_c_*(0) is the maximum value of *N_c_* which is 61. This is the base maximum value and the starting point for all comparisons. Second, we will define *N_c_*(1) which is the expected value of *N*_c_ if only genome or local gene processes that determine overall G+C content are the sole forces shaping codon usage. Codons are used at random with preference towards combinations that equal the G+C content of the overall gene or genome.

**Table 1.** Variations of effective codon number.

| $N_c(M)$ | Effective level of organization | Codon bias constraints | Selective neutrality amongst synonymous codons |
| --- | --- | --- | --- |
| $N_c(0)$ | Base random case | No constraints | All synonymous codons are selectively equivalent. |
| $N_c(1)$ | Genome or gene wide processes | Synonymous codon usage must reflect G+C content. | All synonymous codons with the same number of G or C nucleotides are selectively equivalent. |
| $N_c(2)$ | G+C preference in codon positions | Mutational or selection biases for or against G+C within codons. | All synonymous codons with G/C in the same nucleotide positions treated as selectively equivalent. |
| $N_c(3)$ | Preference for specific codons | Synonymous codon usage must reflect actual frequencies. | Selectively equivalent synonymous codons are dictated by evolutionary forces specific to the gene/organism. |
| Wright's $N_c$ | Any level causing synonymous codon bias. | Synonymous codon usage must reflect actual frequencies. | Selectively equivalent synonymous codons are dictated by evolutionary forces specific to the gene/organism. |

Definitions of various effective codon sizes used in the paper.

Third, will be *N_c_*(2) which is based on the relative G+C contents at each of the three base positions in the codons. This measure reflects the effects of mutational biases or selection pressures that drive preference to codons that match the preponderance or lack of G+C bias for each codon position, especially GC(3). The final measure, *N_c_*(3), which will be shown to very closely approximate *N_c_*, incorporates all other processes that drive codon usage bias. Given the first two measures incorporated genome G+C content and possible mutational bias, *N_c_*(3) reflects these as well as selection processes that select for specific codons and probably overwhelmingly reflect the effects of selection on codon bias.

By comparing these measures in different organisms or even across taxonomy groups, a clear picture of the relative drivers of codon usage bias can be demonstrated as well as outliers that rely almost exclusively on genome, mutational, or selection factors for their distribution of codon usage.

### 1.1 Calculating *N_c_*(1)

In order to create a value of the effective number of codons that reflects only genome wide or gene G+C content we assume that the distribution of codons overall is such that their weighted frequency by G+C content equals the measured G+C content. Codons come in four classifications of G+C content where a codon can have zero, one, two, or three G/C nucleotides. Under the model of random usage, except for G+C content, each synonymous codon has an equal probability of selection if it has the same G+C content as another synonymous codon. Likewise, A+T rich synonymous codons are relatively less/more frequent for G+C rich/poor genes or genomes.

To calculate the distribution of codons by G+C content, we will use the assumption of a maximum entropy distribution in the frequency of the four codon classes subject to the constraints of their weighted average meeting the G+C content. Maximum entropy has been used in the past to measure the effect of G+C bias on codon usage [27] but here we will use the maximum entropy distribution to derive a form of the effective codon number rather than a regression analysis.

Maximum entropy is frequently used as a default hypothesis for a distribution of probabilities subject to given constraints. Maximum entropy is often preferred compared to other distributions, for example a uniform distribution, since random forces subject to the constraints can often be assumed to return the most random (maximum entropy) distribution possible. In other words, given the G+C content of coding sequence, if no other mutational or selection forces are active, codon biases may be assumed to be as random as possible while having the required G+C content.

Assume the probability a codon with a G+C content of *n* is represented by *p_n_* and the overall gene or genome content is *p_GC_*. We thus need to calculate a maximum entropy distribution amongst *p*_0_, *p*_1_, *p*_2_, and *p*_3_ subject to the constraints

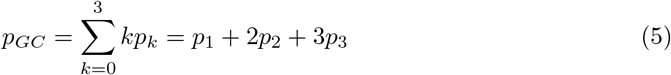

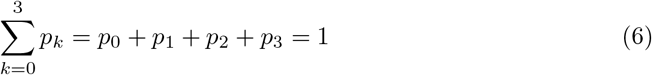

The method of analytically deriving the maximum entropy distribution with the technique of Lagrange multipliers is well studied [28] but for purposes of brevity, this problem reduces to one where it is essential to numerically solve the real root of the order three polynomial

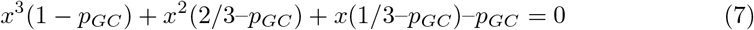

In the equation above *x* = 2^-*λ*/3^ where *λ* is the Lagrange multiplier. Once solved, the individual *p_n_* can be calculated.

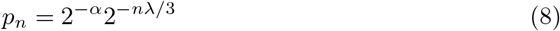

The constant *α* = log_2_ (1 + 2^-*λ*/3^ + 2^-*λ*2/3^ + 2^-*λ*^)

Once we solve for the *p_n_* we can first estimate the relative proportion of synonymous codons based on their G+C values. For example, in the standard code leucine uses codons TTA, TTG, CTT, CTC, CTA, and CTG. These can be arranged into *p*_0_ (TTA), *p*_1_ (TTG, CTT, CTA) and *p*_2_ (CTC, CTG) codons. The codons in each group will be used based on the prevalence of the codon type.

For all codon classes with zero to three codons being G/C, if the G+C content is 50%, all codon classes would be used equally (0.25 × (0 + 1/3 + 2/3 + 1)). However, for example where G+C is 65% the values are *p_0_* = 0.13, *p*_1_ = 0.19, *p*_2_ = 0.28, and *p*_3_ = 0.40. To analyze codon usages by amino acid, these probabilities are equally divided amongst the codons in the type where codons with zero or three G+C bases have 8 combinations while those with one or two have 24 combinations. Therefore the probability of each codon with 0, 1, or 2 G+C bases is 0.016, 0.008, and 0.011. The total probabilities of all six codons for leucine is 0.016 + 3 × 0.008 + 2 × 0.011 = 0.0634 and divide the probability of each to get the probability of each codon representing leucine to be TTA (25%), TTG/CTT/CTA each 13% and CTC or CTG 18%.

Further, we can calculate *N_c_*(1) based on the methodology of Eq. 2. Calculating *H_max_* as the entropy of the distribution of *p_n_*

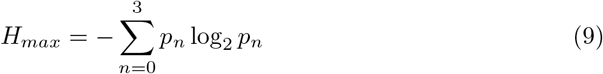

The expected number of codons per G+C type is 2*^H_max_^* though the actual amount can vary due to G+C requirements. Next we multiply this by the expected number of codons per category of 16. This value is the average of the number of codons per category by G+C content. For those codons with G+C of zero or three there only 8 possible combinations while those of G+C of one or two have 24 possible combinations. This averages to 16 and is used as a factor to multiply times the expected number of codons per category. *N_c_*(1) is then defined as

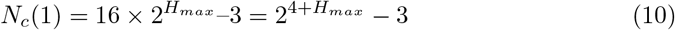

The subtraction of three at the end is to remove the three stop codons in the standard code that are inherent in the assumptions of the calculation of *N_c_*(1). If all codons are equally likely despite G+C content where G+C=50%, *H_max_* = 2 and *N_c_*(1) = 61. This value is the expected value of *N_c_* for random codon usage accounting for genome, or gene, G+C content. The value of *N_c_*(1) is usually not very different from the maximum value of 61 across the common G+C content range of most genes or genomes but as the G+C content becomes increasingly skewed, *N_c_*(1) rapidly decreases. *N_c_*(1) is also symmetric having the same value for genomes of the same G+C or A+T content. Table 2 and Fig 1 demonstrate values of *N_c_*(1) and their trends based on G+C content. It seems for a lower bound of *N_c_*(1) being 20, the minimum and maximum possible G+C content is less than 10% and greater than 90% but due to uneven ratios of G+C across synonymous codons for each amino acid, the bounds are much higher/lower in practice.

**Table 2.** N_c_(1) for various levels of genome G+C content.

| G+C content values | $N_c(1)$ |
| --- | --- |
| 10% / 90% | 29 |
| 20% / 80% | 42 |
| 30% / 70% | 52 |
| 40% / 60% | 59 |
| 50% | 61 |

Expected values of $N_c(1)$ , rounded to the nearest integer, for various values of G+C content.

**Fig 1.**
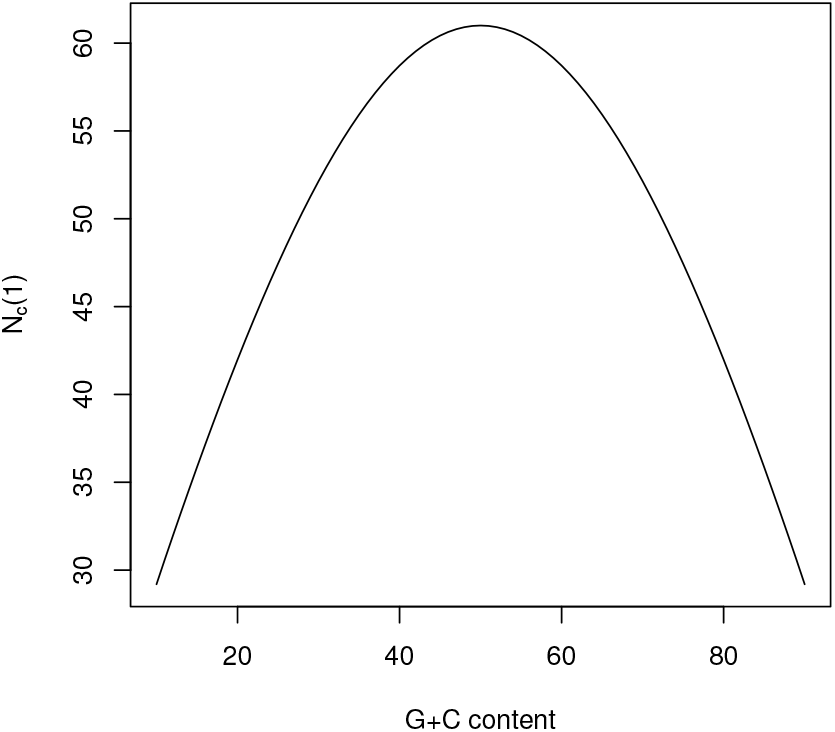
Plot of the expected value of *N_c_*(1) based on genome G+C content.

A close approximation of *N_c_*(1) is given by the equation below, with the variable *GC* as the decimal of the G+C content in range [0,1]

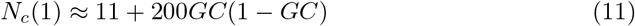

### 1.2 Calculating *N_c_*(2)

Following the calculation of *N_c_*(1) which takes genome wide processes into account, the next level of detail comes from G+C content at the three individual positions within codons which affects codon usage and distribution [29]. The position G+C content, especially GC(3), is theorized to be driven primarily by mutational biases in favor of G+C [13,30–32]. To calculate *N_c_*(2) we will take a simpler route than with *N_c_*(1) while retaining some assumptions.

The entropy content of a single codon position is defined amongst the four nucleotides assuming that synonymous codons with G+C in the same positions will be represented with equal frequency. The entropy at any of the three codon positions *GC*(*N*) can be stated using the frequency *P*_*GC*(*N*)_.

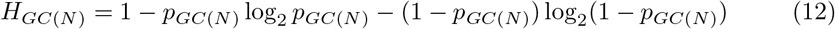

There is a maximum of four if *P*_*GC*(*N*)_ = 50%. The total value of *N_c_*(2) is determined by taking the product of combinations at each three position and removing the three stop codons.

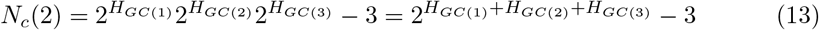

Again the maximum value if *p*_*GC*(1)_ = *p*_*GC*(2)_ = *p*_*GC*(3)_ = 50% is *N_c_*(2) = 61. Because the average of all three positions must equal the total G+C value, the sum of the entropies cannot exceed 4 + *H_max_* and the value of *N_c_*(2) ≤ *N_c_*(1). Given the forces that determine G+C content at each position are largely mutational, *N_c_*(2) is a reflection on the effective number of codons given both G+C content within the genome and mutational forces shaping the G+C content within codons. It does not categorically exclude selection, however, the selection it accounts for are selective forces that only select for/against codons based on the G+C positioning within a codon. Different synonymous codons with G+C at the same positions are considered selectively neutral in terms of *N_c_*(2). The value of *N_c_*(2) is often substantially lower than *N_c_*(1) and is the first reflection of evolutionary forces reducing the effective number of codons towards the value of *N_c_*. The fraction *N_c_*(2)/*N_c_*(1) is a way to normalize the decrease in effective codon size due to forces independent of overall G+C content to compare the relative strength of sequence G+C content and codon nucleotide positional G+C content. Given the latter is often believed to be related to mutation, *N_c_*(2)/*N_c_*(1) can be a method of analyzing these effects normalized for the G+C content of the gene or overall genome.

### 1.3 Calculating *N_c_*(3) and comparison to Wright’s *N_c_*

The final measure of *N_c_* closely approximates the value of *N_c_*. The value *N_c_*(3) takes into account all aspects of codon usage distribution by calculating the total entropy of all sense codons (61 for the standard code though more for others). Accounting for codon usage at the level of the individual codon accounts for almost all information in codon usage bias and is why this closely approximates the traditional *N_c_* value. The sense codon entropy, *H_c_* for the standard code (NCBI codon table 1) is calculated as

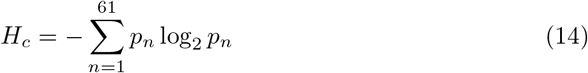

The frequency of the nth codon is represented by *p_n_*. Finally we have

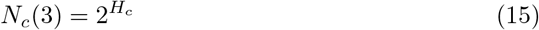

The method of obtaining the effective number of codons is similar to the method of Jost [34] in calculating the effective number of species based on the diversity of species in an area. Subtracting the three stop codons is unnecessary since only the sense codons are accounted for in the calculation. The correspondence between *N_c_*(3) and Wright’s *N_c_* is shown graphically in Fig 2 for a variety of different organisms and Fig 3 for the genes of *Acetobacter pasteurianus*.

**Fig 2.**
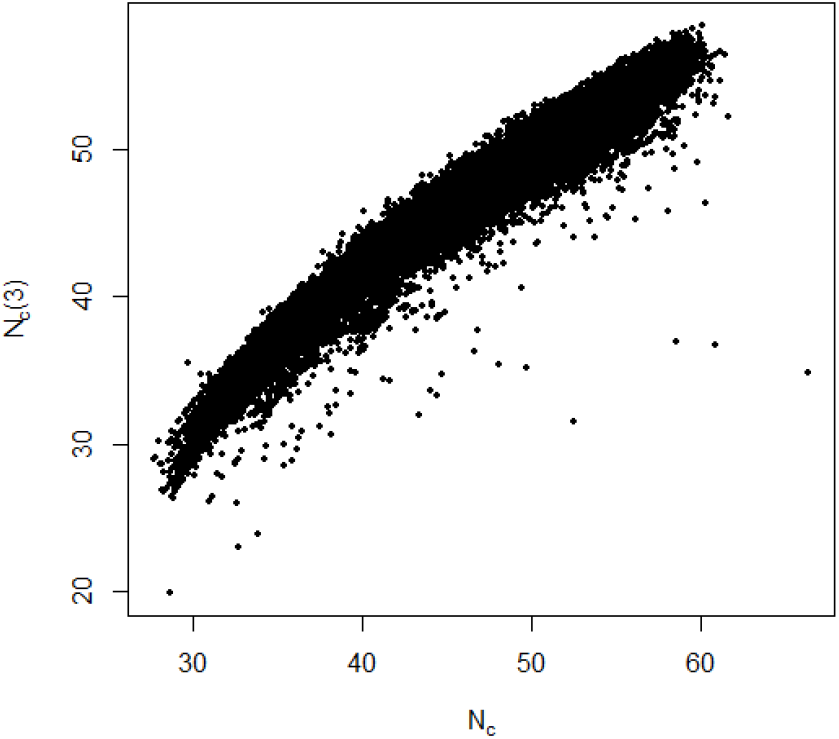
Scatterplot of *N_c_*(3) versus *N_c_* for *N* = 48, 650 genomes. *R*^2^ = 0.94. CDS data obtained from HIVE-CUT RefSeq CDS [33]

**Fig 3.**
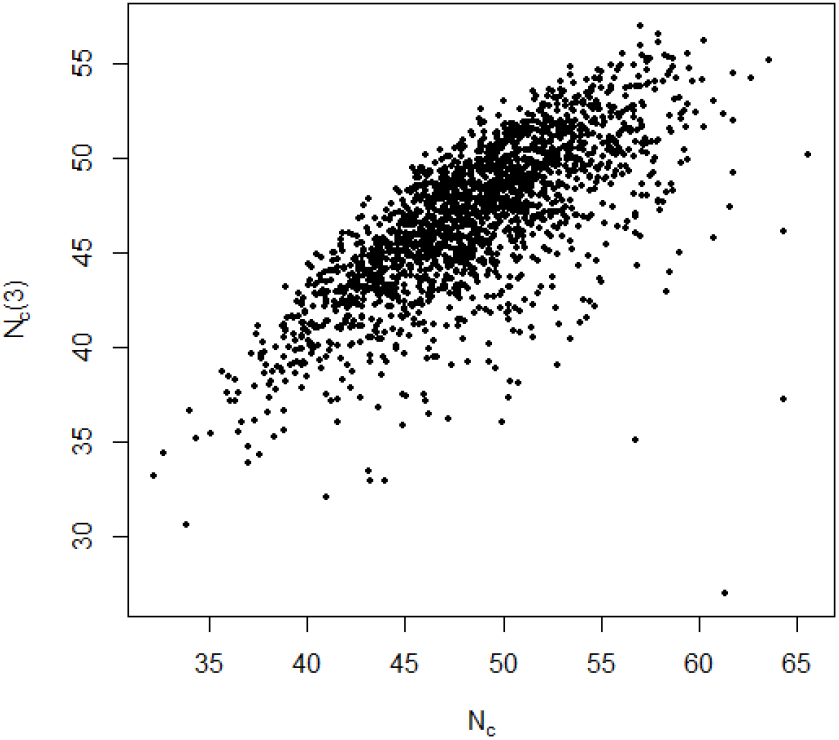
Scatterplot of *N_c_*(3) versus Wright’s *N_c_* for all CDS with at least 200 amino acids for *Acetobacter pasteurianus*, RefSeq genome GCF 000723785.2. *N* = 1, 905 and *R*^2^ = 0.53

There is a close correspondence which is roughly linear at a *R*^2^ = 0.94 in Fig 2. There are some deviations though, typically when a small group of codons have an extremely high frequency as in some viruses or simple eukaryotes, *N_c_*(3) can underestimate *N_c_*. *N_c_*(3) accounts for the balance of forces affecting codon usage bias, most prominently selection or drift which lead to specific synonymous codons being preferred for factors beyond the G+C content overall or mutational biases. In addition, it has the ease of calculation without the necessity of partitioning codons by amino acid as in calculating *N_c_* and other codon usage metrics.

Like *N_c_*(2)/*N_c_*(1) reflected the normalized codon bias due to codon nucleotide effects, *N_c_*(3)/*N_c_*(2) demonstrates the overall codon bias due to codon specific effects by selection or drift that establish preferred codons. The comparison of the two can help understand how different forces shape codon usage bias.

#### 1.3.1 *N_c_*(2) does not reflect codon specific selection

To support the theory that *N_c_*(2) is primarily reflective of mutational or limited selection biases and not selection of individual codons, there are two major details.

First, *N_c_*(2) reflects primarily the effects of the GC(3) content. As predicted in [13], codon usage bias caused largely by patterns in synonymous mutation would be reflected in a relationship between *N_c_* and GC(3) which was approximated as

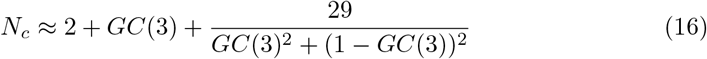

In this equation, GC(3) is in the range [0, 1]. Plots of *N_c_* versus GC(3) are known as *N_c_* plots where the curve in Eq. 16 is shown versus plots of data for different genes or organisms. In Fig 4 *N_c_* plots using *N_c_*(2) and *N_c_*(3) are shown. It is clear *N_c_*(2) closely matches the theoretical curve while *N_c_*(3) is below the curve as is expected when selection lowers the effective number of codons from that bias due only to mutation.

**Fig 4.**
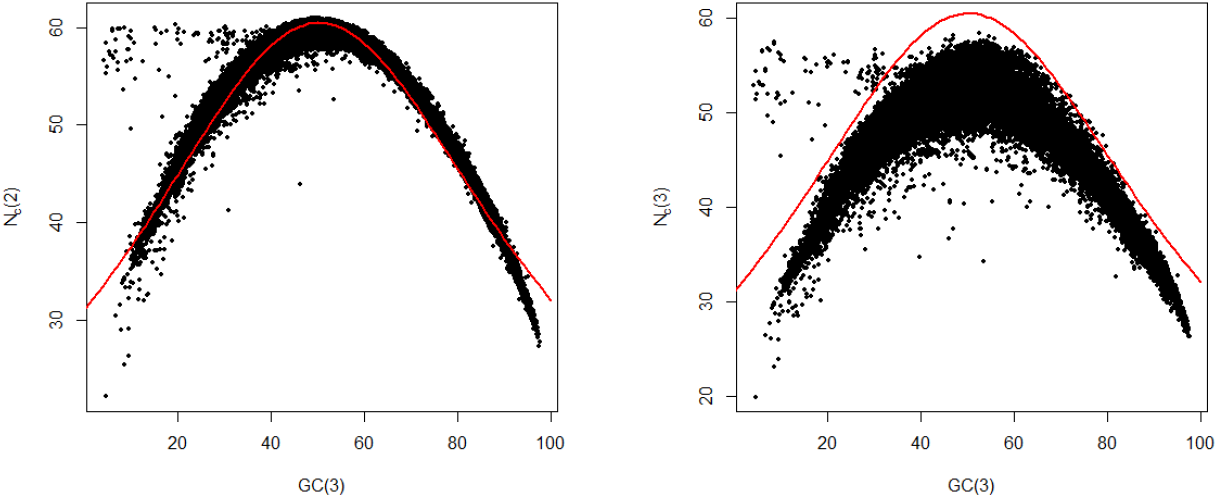
Plots of *N_c_*(2) and *N_c_*(3) versus GC(3) for *N* = 48, 650 organisms from the HIVE-CUT RefSeq database The red line indicates the theoretical value from Eq. 16.

To test the assumption that selective codon usage and not G+C bias at any of the three codon positions drove the value of *N_c_*(3), a numerical simulation was performed across values of G+C from 40% to 75% at steps of 5%. At each G+C content, different values of GC(1), GC(2), and GC(3) were simulated ranging from minimum values of GC minus 20% to maximum values of GC plus 20% at each position in steps of 5% as well. From these 10,000 binary codons (with G or C giving ‘1’ and A or T giving ‘0’) were created to model the codon bias. The frequency of each binary codon was divided by eight to account for all possibilities and *H_c_* and *N_c_*(3) were calculated. As shown in Fig 5, where the line is the average of *N_c_*(3)/*N_c_*(2) and the error bars show the minimum and maximum values, the values of *N_c_*(3) are usually exactly identical to *N_c_*(2) when only the GC(1), GC(2), and GC(3) site contents are considered. Therefore values of *N_c_*(3) substantially lower than *N_c_*(2) are almost surely indicative of selective usage of specific codons.

**Fig 5.**
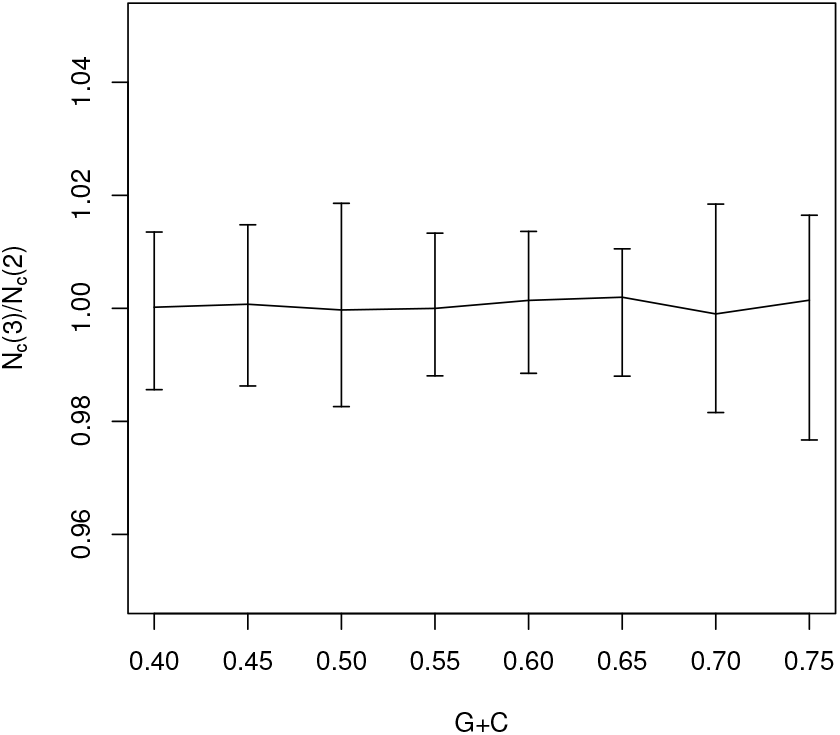
Plot of the average simulated ratio *N_c_*(3)/*N_c_*(2) across multiple values of G+C. Line is the average ratio of the two values while the error bars show the minimum and maximum ratios for each G+C group.

## Results

### 1.4 Using *N_c_*(1), *N_c_*(2), and *N_c_*(3) to understand codon usage bias across organisms

The absolute and relative values amongst the different types of *N_c_*(*M*) can be applied to individual or groups of organisms to investigate factors causing codon usage bias. Using the HIVE-CUT codon usage database [33], codon usage for the CDS from sequenced organisms in RefSeq was analyzed to calculate the various types of *N_c_*(*M*). Differing from HIVE-CUT, *N_c_* was calculated without including stop codons. Only one sequence per taxon ID was used in order to minimize sample bias due to organisms with large numbers of sequences, particularly pathogenic bacteria. In addition, virus betasatellite partial sequences were removed. First using the example of absolute values, eight distinct organisms are compared with all values of *N_c_*(*M*) and *N_c_* in Fig 6.

**Fig 6.**
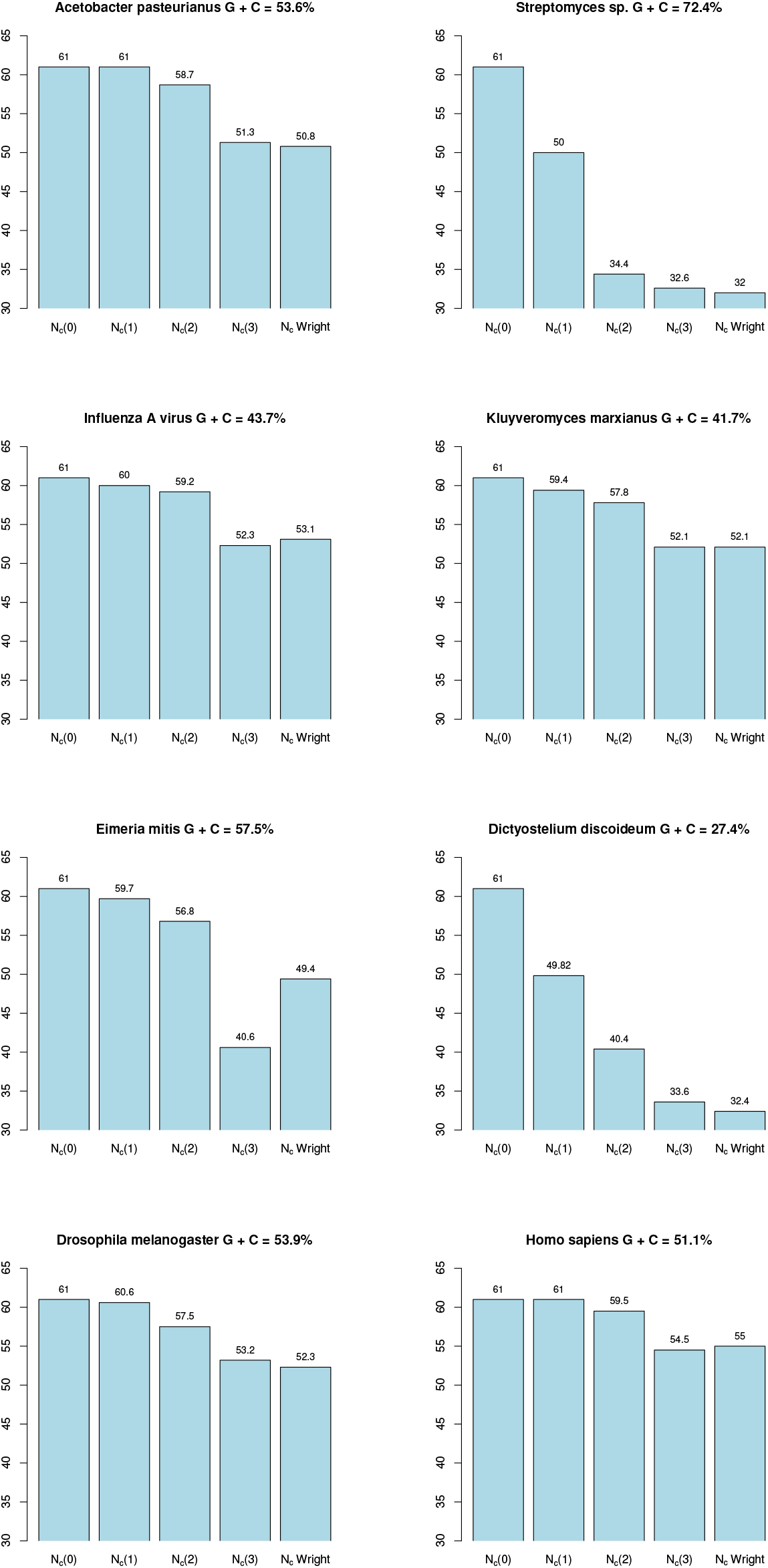
Comparisons of the various effective codon sizes for eight different organisms based on all CDS: *Acetobacter pasteurianus* (Alphaproteobacteria; vinegar fermenting bacterium), *Streptomyces CNT-302* (Actinobacteria), Influenza A (virus), *Kluyveromyces marxianus* (lactose fermenting yeast), *Eimeria mitis* (parasitic protozoan in chickens), *Dictyostelium discoideum* (slime mold), *Drosophila melanogaster* (fruit fly), and *Homo sapiens*. The top of each plot shows the G+C content above the decreasing values of *N_c_*.

Different organisms show relatively different factors influencing their codon bias. For example, in human genomes overall mutation seems to have relatively little effect reducing the effective codon usage only by one from the maximum. *N_c_*(3) and *N_c_* however show a marked decrease to the values of about 55 suggesting selection likely plays a larger, though overall modest, part compared to mutation. More extreme examples are often seen in unicellular organisms and viruses. *Streptomyces* has a large drop from *N_c_*(1) of 50 to a *N_c_*(2) of 34. The difference between *N_c_*(2) and *N_c_*(3) is more moderate down to 32 indicating mutational biases likely drive most of the codon bias, a conclusion identical to that in [39]. An opposite story seems to be the case for the chicken protozoan parasite *Eimeria mitis*. Its *N_c_*(2) of 57 decreases to 41 for *N_c_*(3). However, much of its codon bias is driven by three codons: CAG, AGC, and CGA which collectively account for 28% of all CDS codons and this likely lowers the *N_c_*(3) substantially compared to the *N_c_* of 49 though this is still a substantial reduction.

More informative than absolute numbers are the relative ratios *N_c_*(2)/*N_c_*(1) and *N_c_*(3)/*N_c_*(2). These two ratios normalize the relative difference between effective codon sizes across different organisms in a way absolute numbers cannot. Therefore we can compare individual organisms or even look at wide groups using the first ratio as a measure of the reduction of codon size due to mutational biases while the second is a reduction largely due to specific codon selection pressures. The latter, *N_c_*(3)/*N_c_*(2), can also be interpreted as the deviation of actual codon usage in amino acids from the baseline expected given GC biases in each codon position.

### 1.5 Novembre’s *N_c_*′ comparison

The metric *N_c_*′ of [16], which is an alternate measure of *N_c_* reflecting deviations from expected codon usage rather than even codon usage as in Wright’s *N_c_*, can be approximated by *N_c_*(3)/*N_c_*(2). The derivation of *N_c_*′ is based on using the chi-square deviation of actual and expected codon usage amongst synonymous codons to create a metric similar to the homozygosity used in Wright’s *N_c_*. This metric is then used in an identical manner to the homozygosity in Wright’s *N_c_* to calculate *N_c_*′. The strength of *N_c_*′ lies in allowing for various models for expected codon usage to be tested against actual codon usage to create an effective codon number description.

In our analysis of *N_c_*′, we will adopt an expected codon usage from the codon usage determined by *N_c_*(1) given the genome or CDS G+C content. However, to calculate *N_c_*′ we must determine not only the overall effective codon number but also the expected frequency of each codon. Overall, codon usages are determined according to the GC percentage in each of the three nucleotide positions. If overall G+C content is the only determinant of codon frequency, the default codon usage frequencies for each codon can be calculated by using the number of G/C nucleotides and the probabilities determined in the calculation of *N_c_*(1) (i.e. codons with two G/C nucleotides have probability given by *p*_2_ from equation 8). Given these expected codon frequencies, chi-square is then calculated using these expected frequencies and the actual frequencies for each amino acid. These are then used in calculating *N_c_*′ per [16]. In Fig 7 this is demonstrated by showing a plot of *N_c_*′ versus *N_c_*(3)/*N_c_*(2). While many values of *N_c_*(3)/*N_c_*(2) can be represented by multiple values of *N_c_*′, the overall trend is positive showing a significant correlation between the two measures to detect deviations from expected codon usage given mutational or codon site selection processes.

**Fig 7.**
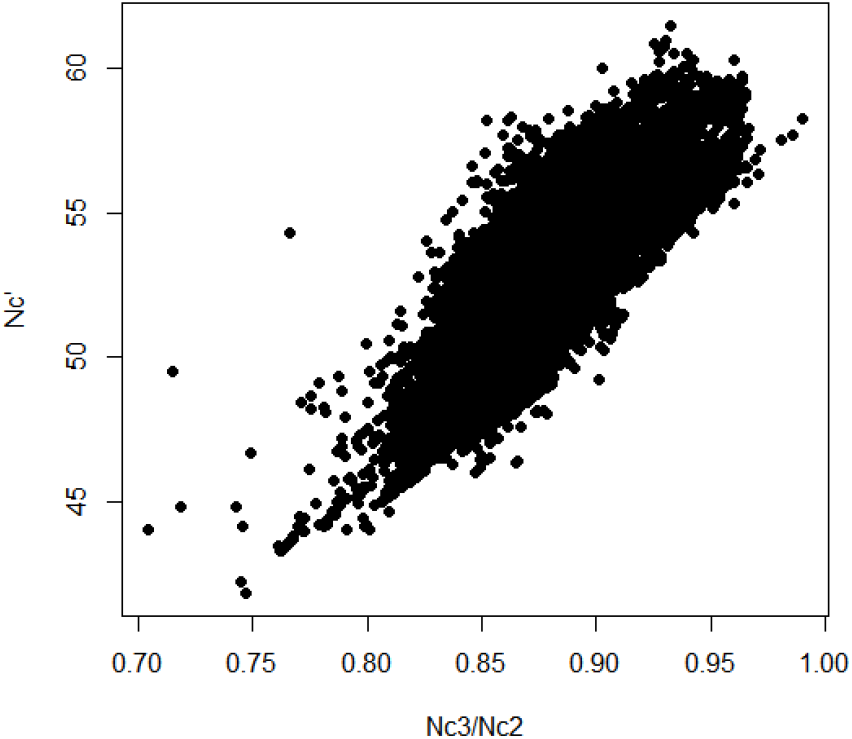
Plot of *N_c_*′ versus the ratio *N_c_*(3)/*N_c_*(2) for *N*=42, 427. *R*^2^=0.53

### 1.6 *N_c_* ratio results across organisms

In Table 3 the organisms from Fig 6 have their ratios listed. Most insightful, however, is a plot of *N_c_*(3)/*N_c_*(2) vs. *N_c_*(2)/*N_c_*(1) for large groups of related organisms. These allow us to see across a wide span of organisms, how patterns of mutational or selection forces shaping codon bias occur. In the plots of Figs. 8, 9, 10, 11, and 12, this is shown for phyla across Bacteria, Archaea, several categories of viruses, the phylum of Chordata, various invertebrate phyla and for various mitochondrial and plant chloroplast sequences.

**Table 3.**
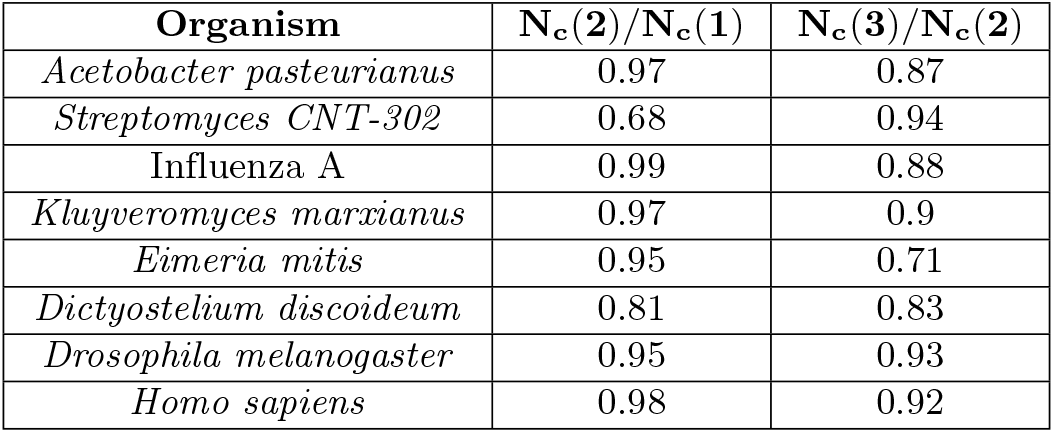
Values of *N_c_*(2)/*N_c_*(1) and *N_c_*(3)/*N_c_*(2) for the eight organisms.

**Fig 8.**
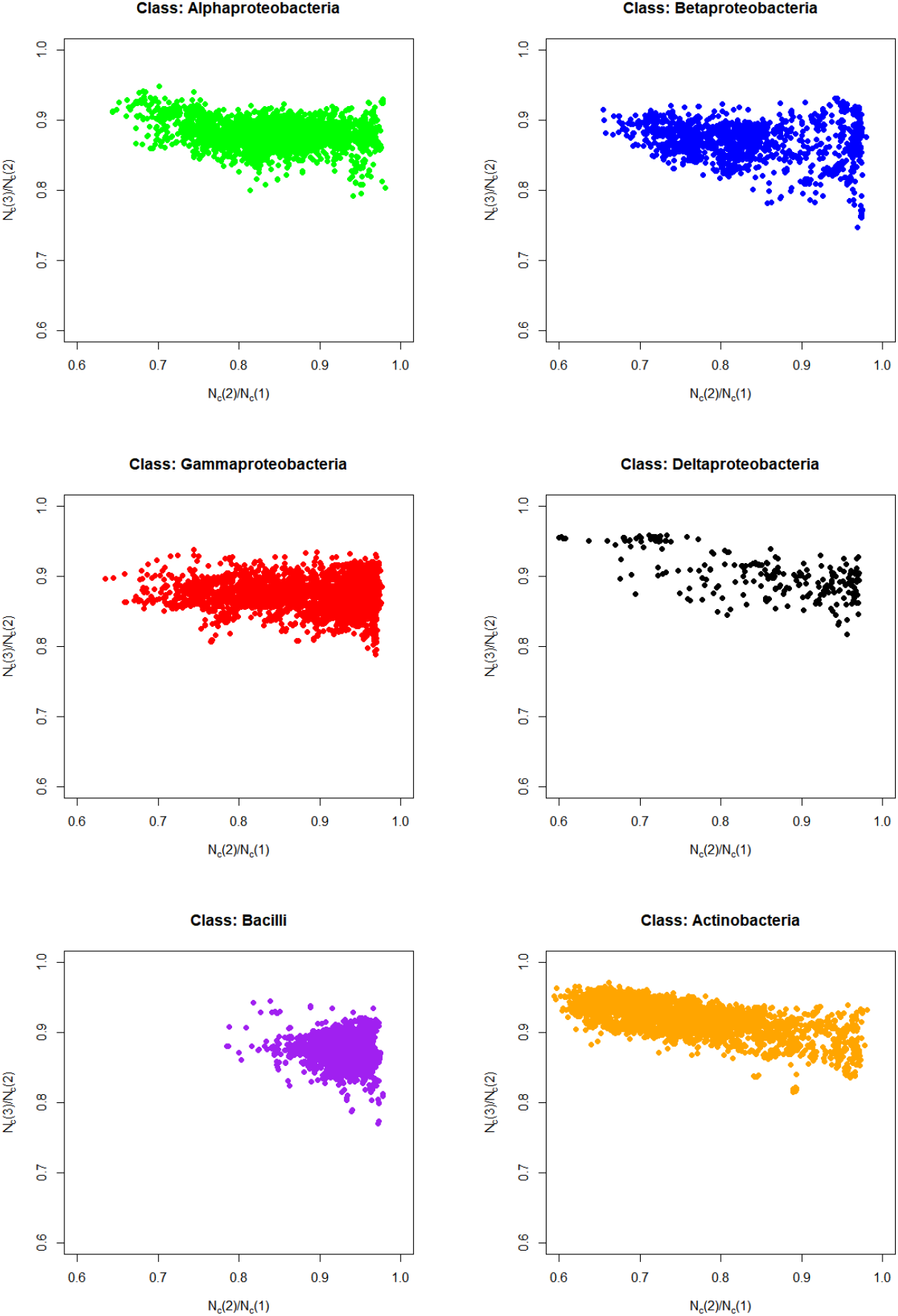
Ratios *N_c_*(3)/*N_c_*(2) versus *N_c_*(2)/*N_c_*(1) across the bacteria classes of Alphaprotebacteria (*N* = 3, 293), Betaproteobacteria (*N* = 2, 114), Gammaproteobacteria (*N* = 11, 437), Deltaproteobacteria (*N* = 235), Bacilli (*N* = 9, 646), and Actinobacteria (*N* = 6, 380). N designates the number of distinct taxon IDs.

**Fig 9.**
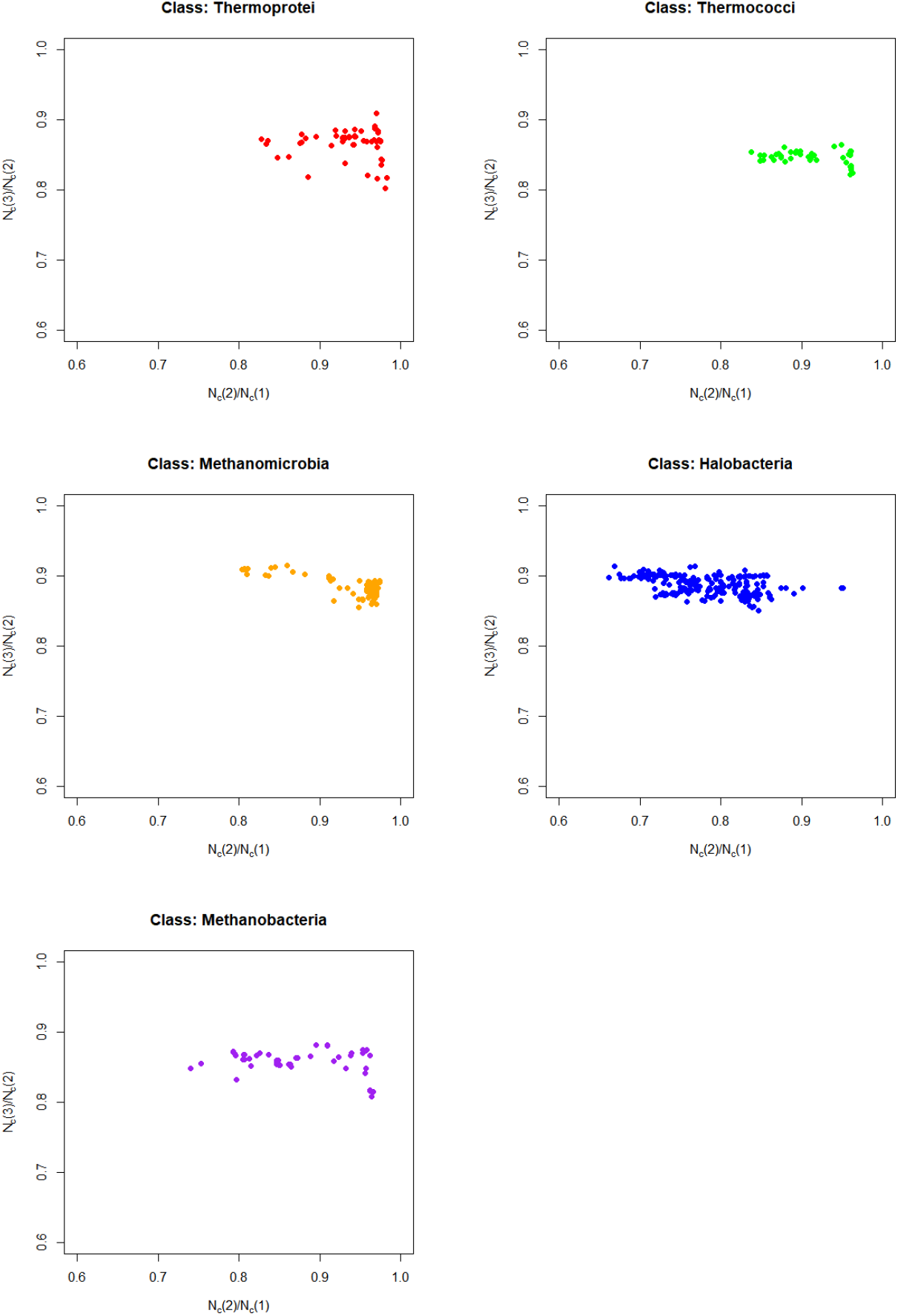
Ratios *N_c_*(3)/*N_c_*(2) versus *N_c_*(2)/*N_c_*(1) across the Archaea classes of Thermoprotei (*N* = 71), Thermococci (*N* = 44), Methanomicrobia (*N* = 85), Halobacteria (*N* = 208), and Methanobacteria (*N* = 67).*N* designates the number of distinct taxon IDs.

**Fig 10.**
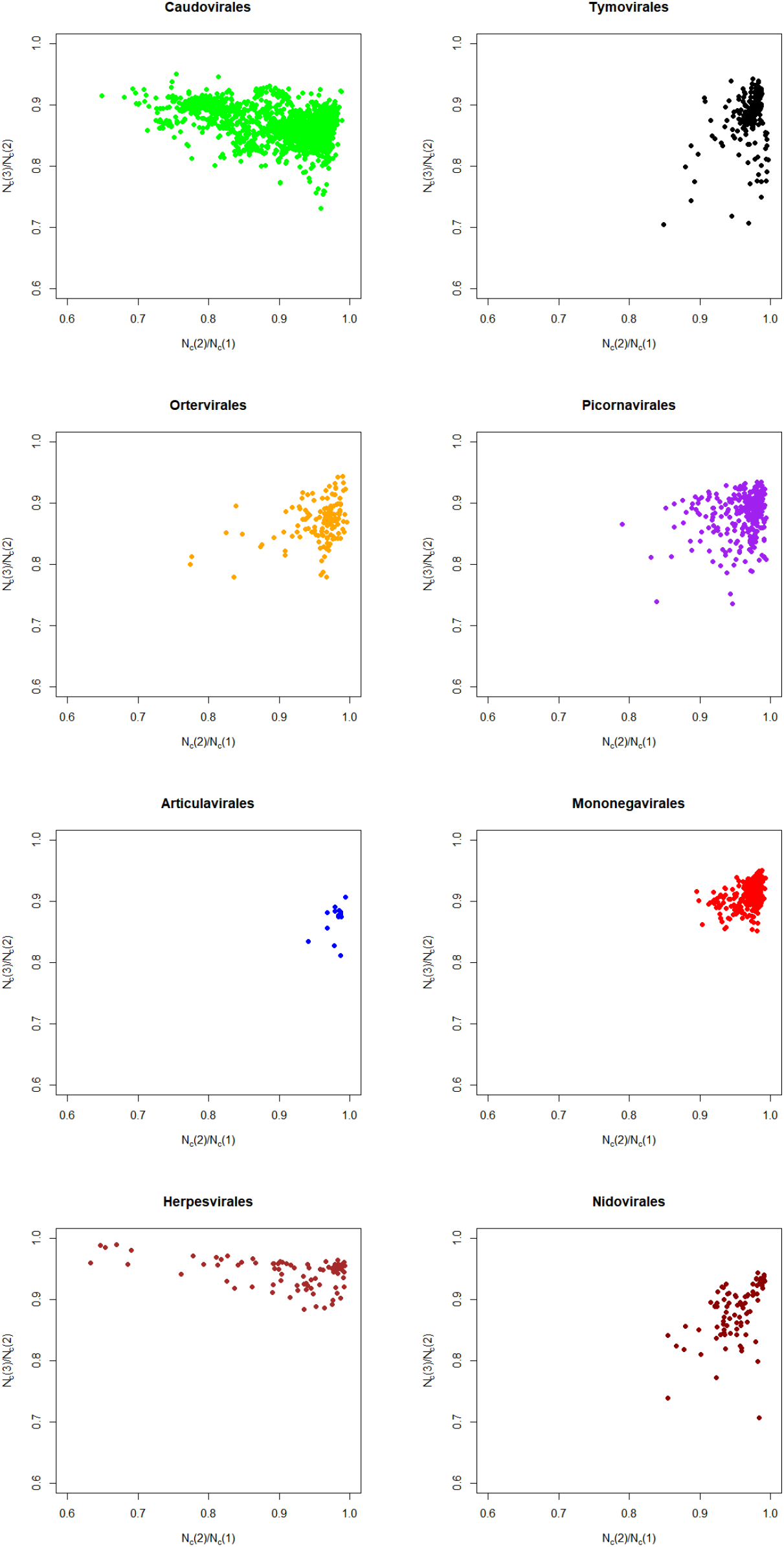
Ratios *N_c_*(3)/*N_c_*(2) versus *N_c_*(2)/*N_c_*(1) across the virus classes Caudovirales (*N* = 2, 049), Tymovirales (*N* = 190), Ortervirales (*N* = 134), Picornavirales (*N* = 282), Articulavirales (*N* = 15), Mononegavirales (*N* = 246), Herpesvirales (*N* = 69), and Nidovirales(*N* = 86). N designates the number of distinct taxon IDs.

**Fig 11.**
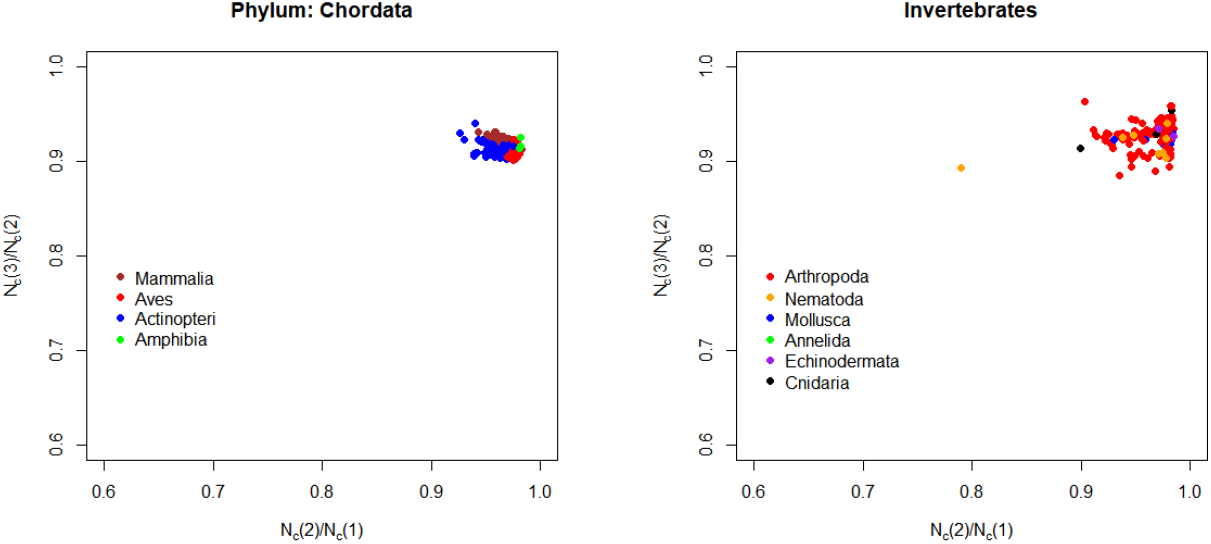
Ratios *N_c_*(3)/*N_c_*(2) versus *N_c_*(2)/*N_c_*(1) across the Phylum Chordata (Mammalia (*N* = 115), Aves (*N* = 62), Actinopteri (*N* = 50), Amphibia (*N* = 3)) and various phyla of invertebrates (Arthropoda (*N* = 126), Nematoda (*N* = 8), Mollusca (*N* = 8), Annelida (*N* = 1), Echinodermata (*N* = 2), Cnidaria (*N* = 6)). N designates the number of distinct taxon IDs.

**Fig 12.**
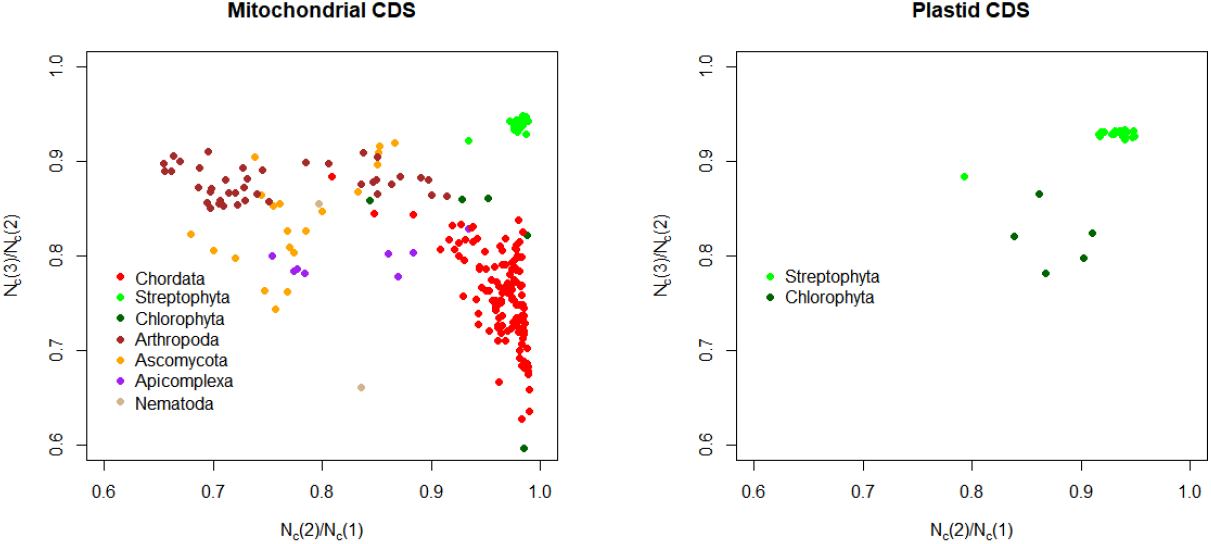
Ratios *N_c_*(3)/*N_c_*(2) versus *N_c_*(2)/*N_c_*(1) for mitochondria (Chordata (*N* = 164), Streptophyta (*N* = 25), Chlorophyta (*N* = 5), Arthropoda (*N* = 40), Ascomycota (*N* = 20), Apicomplexa (*N* = 8), Nematoda (*N* = 2))and chloroplasts (Streptophyta (*N* = 57) and Chlorophyta (*N* = 5)). N designates the number of distinct taxon IDs.

The overall patterns range from the relatively consistent and high ratios for vertebrates to the wide variations of unicellular organisms and mitochondria. In particular, Archaea and Bacteria tend to show a relatively restricted variation in codon bias due to selection but wide variation due to mutational processes with mutational biases with many types of bacteria having relatively high *N_c_*(3)/*N_c_*(2) near one but with much lower *N_c_*(2)/*N_c_*(1) demonstrating the effects of pressures on G+C content in codon positions. Viruses have the widest diversity with either or both mutation and selection playing a large part across many different viruses. While individual organisms may show stronger selection, on balance, selection only seems consistently significantly stronger than mutation in vertebrate mitochondria.

### 1.7 Using *N_c_*(1), *N_c_*(2), and *N_c_*(3) to understand codon usage bias within organisms

The techniques described above can also be used to analyze the codon usage bias across different CDS within a single organism’s genome. The techniques of analysis are basically the same, however, in order to restrict the analyses to those CDS which are least likely to have skewed codon bias due to a short sequence length, the organisms presented here are analyzed using only those CDS which contain at least 200 codons. The results shown in Fig 13 as density plots are consistent and different from the analysis across organisms. For one, biases represented by the decrease of the ratio *N_c_*(2)/*N_c_*(1) seem dominant in the variation codon usage biases across organisms, while biases in codon usage due to selection of specific codons, represented by *N_c_*(3)/*N_c_*(2) seems to dominate the variation of codon usage bias within genomes. This may not be unexpected since within a single organism, processes that create mutational or non-codon specific selection biases are likely rather uniform while genes that require high expression are more likely to have biases in the content of their codons in order to maximize efficiency.

**Fig 13.**
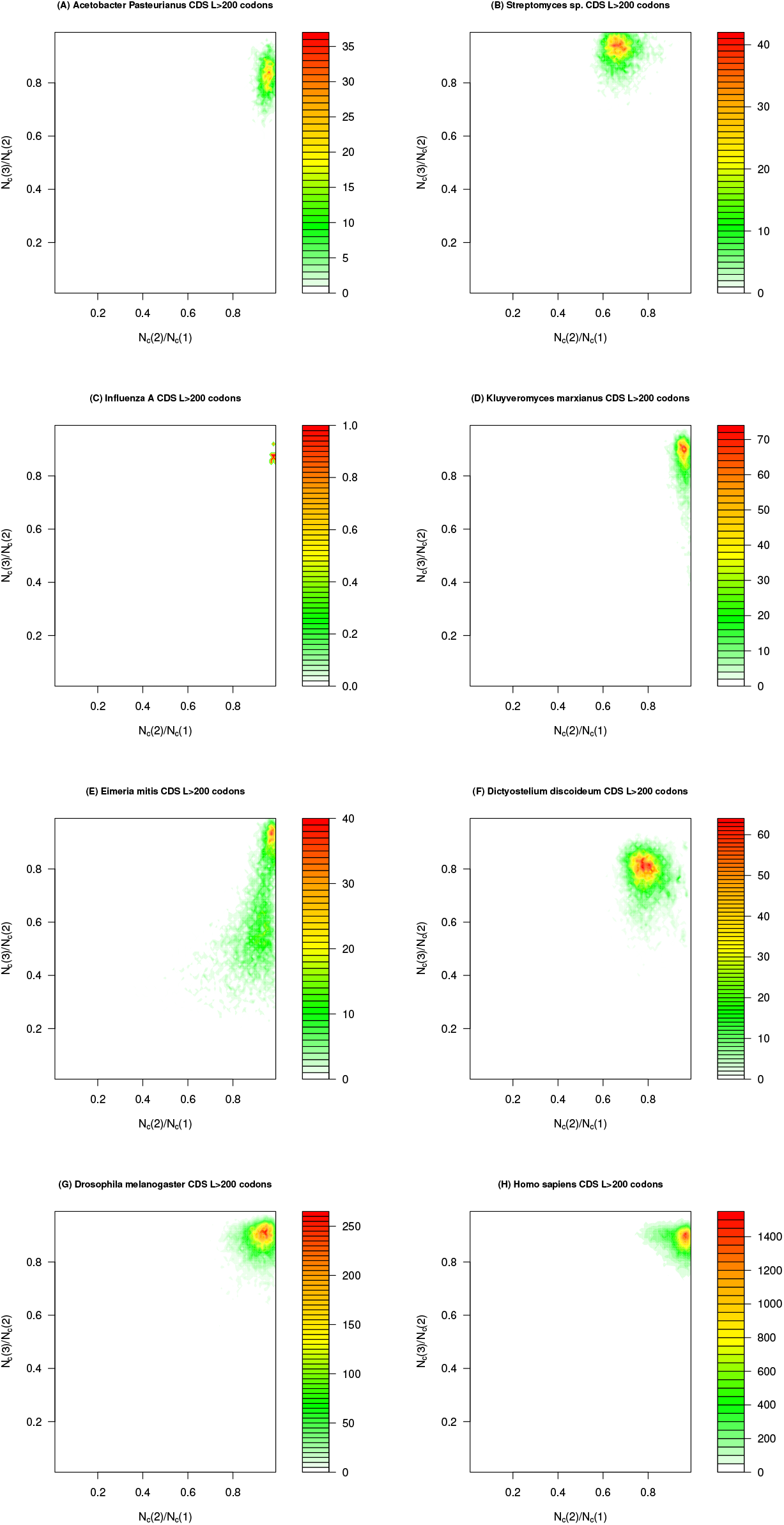
Density plots of the CDS counts for various combinations of the ratios *N_c_*(3)/*N_c_*(2) versus *N_c_*(2)/*N_c_*(1) within various organisms for CDS with at least 200 codons. (A) *A. pasteurianus N* = 1, 905, (B) *Streptomyces sp. CNT-302 N* = 4, 248, (C) *Influenza A N* = 9, (D) *K. marxianus N* = 4, 205, (E) *E. mitis N* = 6, 114, (F) *D. discoideum N* = 9, 937, (G) *D. melanogaster N* = 25, 236, and (H) *H. sapiens N* = 105, 072.

Like variation across organisms, simpler organisms such as bacteria, yeast, or protozoa seem to manifest more variation within the genome while complex multicellular organisms show such variation in a more restricted range.

## Conclusion

The methods and results in this paper formalize the distinction of the forces operating on codon usage bias at all levels. While two of the metrics *N_c_*(2) and *N_c_*(3) closely approximate earlier metrics from [13] for *N_c_* assuming only mutation and *N_c_* overall, they possess some distinct advantages. First, *N_c_*(2) directly incorporates G+C content at all three sites though GC(3) is usually decisive. This can give more accurate results when GC(3) is extremely skewed high or low and the approximation in Eq. 16 overestimates the effective codon number. While similar to *N_c_*, *N_c_*(3) is much easier to calculate requiring only the frequency of each codon and the number of stop codons. The knowledge of the number of degenerate codons per amino acid and adjustments for situations where *N_c_* needs to account for unused or heavily skewed codon usages are unnecessary. The measure *N_c_*(1) is the first measure to genuinely give a base random case for codons incorporating G+C content and not requiring the assumption of equal usage.

The new metrics also allow for tentative testing of the likelihood codon usage in organisms or genes is driven by G+C content, mutation, or selection. Comparisons across organisms can give insight into overall processes that generate codon bias that are not illuminated by one number alone. For example, genes or isochores with similar values of *N_c_*(2)/*N_c_*(1) possibly have similar processes affecting the choice of nucleotides at codon positions, often caused by mutation, though the G+C content differences between them give different values of *N_c_*. Similarly, organisms or genes with similar values of *N_c_*(3)/*N_c_*(2) can have similar biased selection of synonymous codons though they are not necessarily the same codons being selected in each organism. Differences in these amongst relatively related organisms can give clues to possible evolutionary events that can be investigated further.

A prominent example of this is the low value of *N_c_*(3)/*N_c_*(2) (0.71) for *Eimeria mitis* and other members of the *Eimeria* genus. The *Eimeria* genus are coccidia parasites that infect a wide variety of animals but are best known for causing major economic losses for chickens and other domesticated fowl. However, they have a fascinating and unique organization of their genomes that greatly impacts codon bias [35–38]. In particular, the coding sequences are replete with repeats of the codon CAG, and its permutations, primarily coding for the amino acids alanine and glutamine. In [36] it was found that CAG and its permutations were part of runs of amino acids at least seven residues long in 57% of the genes of *Eimeria tenella* with an average of 4.3 copies of CAG per gene. The selection of CAG and permutations such as TGC, GCA, AGC, and CTG account for a huge portion of the genome and makes *Eimeria* unique amongst eukaryotic organisms in this type of organization. This selection of these specific codons drives down *N_c_*(3) and the ratio *N_c_*(3)/*N_c_*(2) and was clearly apparent from the data without details of *Eimeria* known beforehand. While most organisms do not have such extreme ratios, this demonstrates their utility in analysis.

Comparing across organisms and within genomes as shown in the figures seems to show a pertinent pattern despite a few exceptions. First, the differences amongst organisms can be wide but large variations are more often driven by different biases in codon nucleotides shown by *N_c_*(2)/*N_c_*(1) versus the different levels of codon selection shown by *N_c_*(3)/*N_c_*(2). This is especially true in unicellular organisms though some groups of viruses show widely different selection pressures on overall codon usage. On the other hand large variations in specific synonymous codon usage preference amongst genes within organisms seem driven by selection which is consistent with the observation codon usage bias often varies with genes based on frequency of expression.

Many of the results at the organism or gene level also corroborate previous theories about the roles of various evolutionary forces on codon usage bias. As stated earlier, *Streptomyces sp*. shows codon usage likely largely shaped by mutation [39] with relatively muted influence of codon selection both at the whole chromosome as well as the CDS level. On the other hand, the codons in the Influenza virus show little influence of mutation but the definite marks of codon selection [40] with one of the highest *N_c_*(3)/*N_c_*(2) being the codons of the surface protein hemagglutinin with a *N_c_*(2)/*N_c_*(1) of 0.97 and *N_c_*(3)/*N_c_*(2) of 0.85.

Another closely corresponding result for within genome comparison is the codon bias of plant chloroplast genes. In particular, Morton and others [41–43] have noted the atypical codon bias of *psbA* and how it may have been shaped by selective forces though such forces are possibly ancestral and now relaxed [43]. The relative ranking of other genes also closely matches those found by CAI in [42]. A plot of these genes is shown in Fig 14

**Fig 14.**
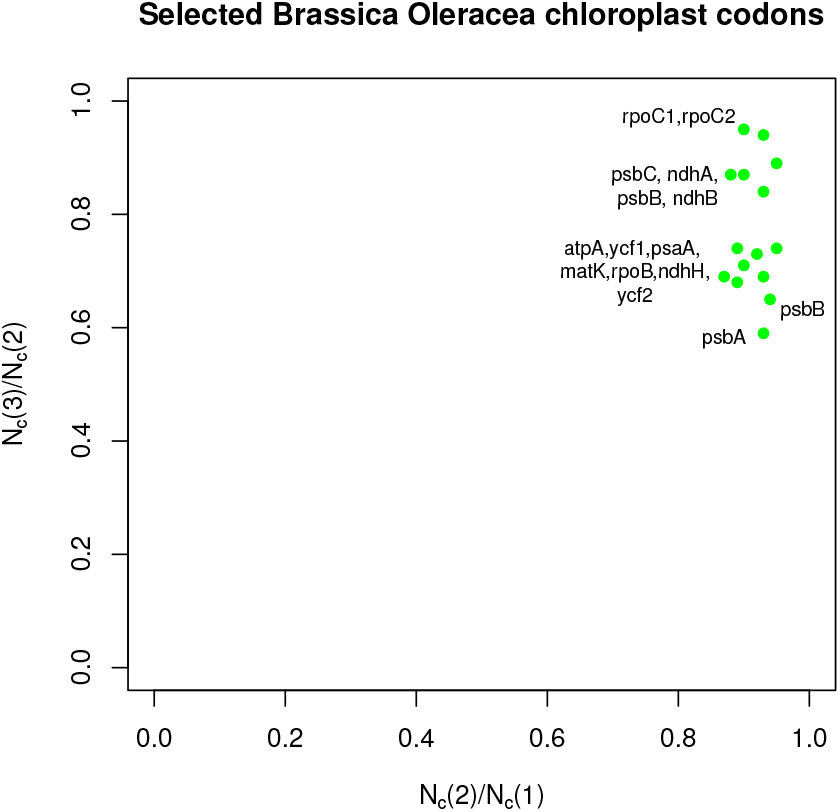
Scatterplot of *N_c_*(3)/*N_c_*(2) versus *N_c_*(2)/*N_c_*(1) for selected chloroplast genes in *Brassica oleracea* isolate RC34 Genbank MG717288.1.

Comparing genes within organisms, where mutational biases are relatively constant, shows that selection, driven by various efficiencies or adaptations, drives most of the differentiation in codon usage bias. Therefore it seems broadly that the values of *N_c_*(2)/*N_c_*(1) show wider variation among organisms and *N_c_*(3)/*N_c_*(2) show wider variation within organisms.

The overall theory underlying the methods in this paper is that each force biasing codon usage, from the genome or gene level to the mutational and selective processes, drives a reduction in the effective codon size from its theoretical maximum of 61 to the final value of *N_c_*(3). Analyzing each of these separately is possible using information theoretic methods applied to combinatorics without making unreasonable or unrealistic assumptions about the underlying genetic mechanisms. The relative amount of reduction in the effective codon number between each analysis is a generalizable and comparable across or within organisms to investigate the causes of codon usage bias despite differences in genome G+C content or codon site G+C content.

Finally, by allowing the causes of codon usage bias to be compared across wide groups of organisms, a consistent study of the causes of codon bias in homologous sequences compared across phylogenetic trees can perhaps give more clues to evolutionary processes and relations amongst organisms. As always, detailed work at the organism level is essential to unveiling the details.

## Acknowledgments

I would like to thank Dr. Hiroyuki Arai for helpful data on *A. Pasteurianus*.

